# Best practices for analyzing imputed genotypes from low-pass sequencing in dogs

**DOI:** 10.1101/2021.04.29.441990

**Authors:** Reuben M. Buckley, Alex C. Harris, Guo-Dong Wang, D. Thad Whitaker, Ya-Ping Zhang, Elaine A. Ostrander

## Abstract

Although DNA array-based approaches for genome wide association studies (GWAS) permit the collection of thousands of low-cost genotypes, it is often at the expense of resolution and completeness, as SNP chip technologies are ultimately limited by SNPs chosen during array development. An alternative low-cost approach is low-pass whole genome sequencing (WGS) followed by imputation. Rather than relying on high levels of genotype confidence at a set of select loci, low-pass WGS and imputation relies on the combined information from millions of randomly sampled low confidence genotypes. To investigate low-pass WGS and imputation in the dog, we assessed accuracy and performance by downsampling 97 high-coverage (>15x) WGS datasets from 51 different breeds to approximately 1x coverage, simulating low-pass WGS. Using a reference panel of 676 dogs from 91 breeds, genotypes were imputed from the downsampled data and compared to a truth set of genotypes generated from high coverage WGS. Using our truth set, we optimized a variant quality filtering strategy that retained approximately 80% of 14M imputed sites and lowered the imputation error rate from 3.0% to 1.5%. Seven million sites remained with a MAF > 5% and an average imputation quality score of 0.95. Finally, we simulated the impact of imputation errors on outcomes for case-control GWAS, where small effect sizes were most impacted and medium to large effect sizes were minorly impacted. These analyses provide best practice guidelines for study design and data post-processing of low-pass WGS imputed genotypes in dogs.

## Introduction

The price per marker for a genotyping assay can have a large influence on the success of genetic association studies. In dogs, DNA genotyping arrays, which provide hundreds of thousands of genotypes at relatively low costs, are highly beneficial for mapping loci (Awano et al. 2009; Hayward et al. 2016), characterizing genetic architecture (Boyko et al. 2010; Friedrich et al. 2019), and defining breed and population structure (Shannon et al. 2015; Ali et al. 2020). However, DNA genotyping arrays are limited by various known and unknown biases that occur during marker selection and probe design that cannot be removed without redesigning a new DNA array, which is an expensive and time-consuming process. An alternative similarly priced approach is low-pass whole genome sequencing (WGS) and imputation (Martin et al. 2021). Rather than assigning genotypes based on high confidence calls across a finite set of loci, low-pass WGS combines information from millions of randomly sampled low confidence variant calls to impute likely genotypes from a reference panel, comprised of a large collection of WGS datasets representing potential haplotypes found within a population. Since low-pass WGS isn’t biased towards sampling specific loci, the only limiting factor is the reference panel used. Therefore, the utility of previous datasets never diminishes.

Due to its flexibility and scalability, low-pass sequencing and imputation has been applied to humans (Rubinacci et al. 2021; Wasik et al. 2021) and other mammalian species (Benjelloun et al. 2019; Piras et al. 2020; Snelling et al. 2020; Nosková et al. 2021). Results in humans demonstrate that low-pass WGS and imputation provide more accurate genotypes than those imputed using array data, leading to increased power for genome wide association studies (GWAS) and more accurate polygenic risk score calculation. Piras et al. (2020) used low-pass WGS and imputation to identify candidate loci for canine idiopathic pulmonary fibrosis in West Highland white terriers (CPSF7 and SDHAF2). While successful in this case, the existing 350 dog breeds present a unique problem for conducting GWAS studies, as the existing structure of each breed, its history, and genome homogeneity are distinct (Ostrander et al. 2017). In the absence of empirical evidence for developing optimal strategies for study design and data processing, the probability of poor performance and misleading results is unknown. As many dog breeds and populations have only been sequenced to low levels, the development of a generalizable set of rules for low pass WGS imputation across breeds would convert much of the existing data from low to high applicability, thus accelerating the dog as a genetic system for studies of canine and human health.

Here, we present an analysis of imputation accuracy of low-pass WGS in the context of canine genomics and establish optimized approaches for study design and data processing. We analyzed imputed genotypes from 97 test samples from 58 different breeds, many of which are not included in the reference panel containing the haplotypes used for imputation. We assessed the impact of minor allele frequency (MAF) on genotyping accuracy and determined whether it was better to use MAFs generated from imputation or to use reference panel MAFs. Finally, we investigate the impact of imputation errors on study design by determining the necessary sample sizes and case-control ratios for a sufficiently powered case-control GWAS.

## Results

### Breed composition of the imputation reference panel and test datasets

The test dataset used for assessing imputation accuracy consists of 97 samples that met selection criteria (Methods). Of the 676 samples used in the reference panel, whose IDs were provided by Gencove, Inc. (New York, NY), 554 were matched to a previously published dataset (**Fig 1A**) (**Supplemental Table S1**) (Plassais et al. 2019). Since a large portion of samples were shared between the Gencove reference panel and the published dataset, we opted to use the publicly available VCF file as a stand-in for the reference panel VCF. The individual breeds comprising the reference samples were compared to the breeds in the Plassais et al. (2019) dataset and the breeds from our test dataset (**Fig 1B**). Only 13 breeds were identified as unique to the published data set and they ranged from one to three members each (**Fig 1C**) (**Supplemental Table S2**). The remaining breeds shared between the reference panel and the published dataset typically contain similar numbers of individuals, with village dogs (VILL), Yorkshire terriers (YORK), wolves (WOLF), Labradors (LAB-), golden retrievers (GOLD), and unknown breeds (UNKN) being the six most popular breed designations in both datasets (**Fig 1C**). Within the test samples, 23 breeds were shared with the reference panel and four breeds were shared with the Plassais et al. (2019) dataset only, while 32 breeds were unique to the test samples (**Fig 1B**). In terms of member frequency per breed, the test samples had no more than four Sealyham terriers, the most common breed within the test samples. In addition, the total members per breed were relatively evenly distributed between breeds unique to the test samples and breeds found in other datasets (**Fig 1C**).

**Fig 1:**
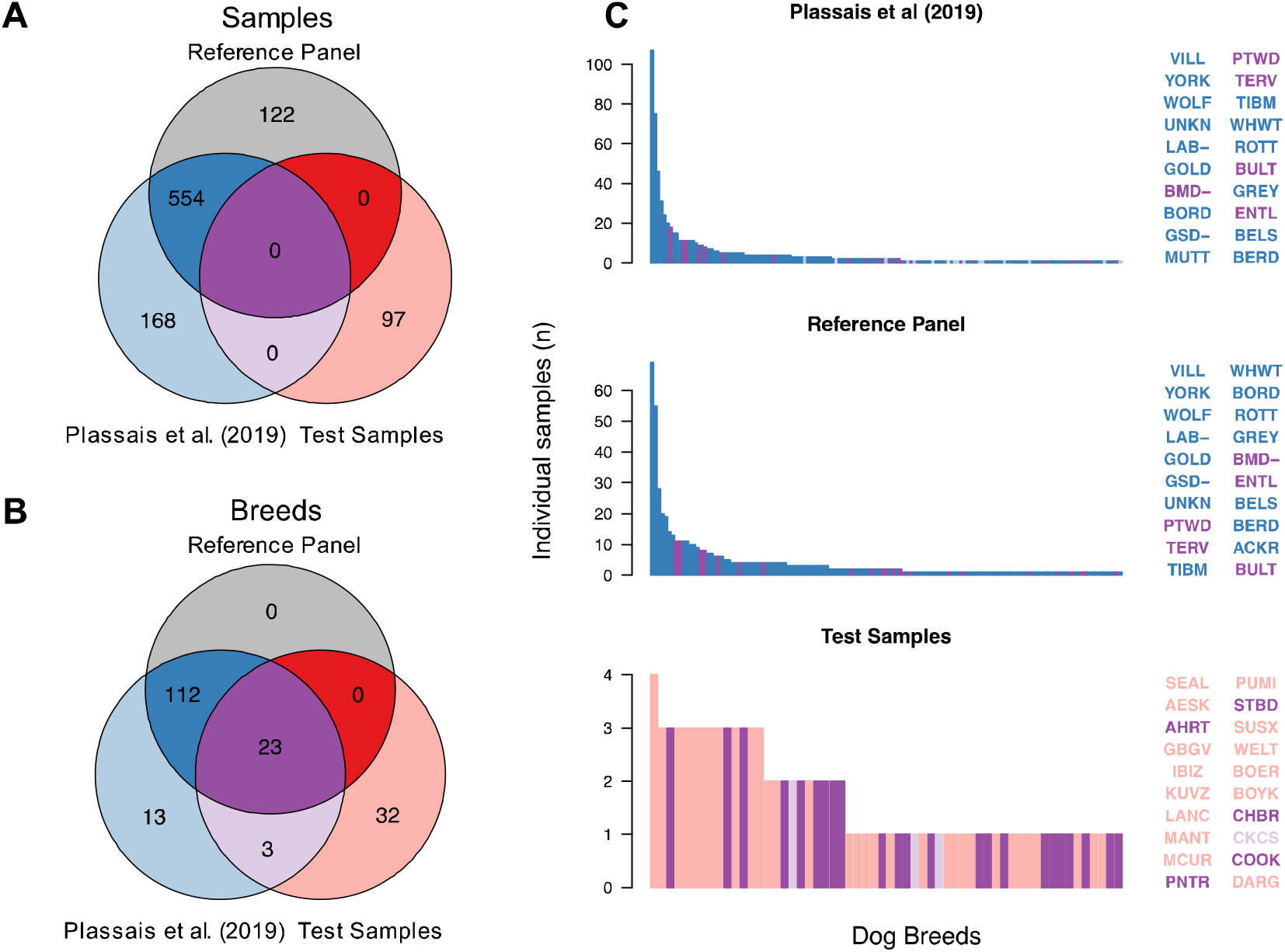
Test samples belong to a wide variety of breeds with most breeds likely not found within the imputation reference panel. **A)** Sample membership within each dataset. Reference panel IDs could not always be linked to a publicly available dataset. **B)** Breed membership among each dataset. Reference panel dogs whose IDs could not be linked to a publicly available sample have no breed label. **C)** Breed frequency across each dataset. Using the colors from the Venn diagram in B, bar colors represent the population a specific breed can be found in. Labels to the left of each bar chart identify the 20 most common breeds. Breeds in bar charts are sorted by most to least common.

To evaluate breed representation at a higher level, we determined the clade membership of all known breeds across the test dataset and the reference panel dataset (Methods) (**Table 1**) (**Supplemental Table S3**). Altogether, the Pinscher and Hungarian clades were unrepresented in the reference panel. This is, perhaps, because each of these clades contains only two breeds (Parker et al. 2017). Other clades of concern due to underrepresentation include the American terrier, Asian toy, small spitz, and toy spitz clades, which have less than three representative samples within the known breeds of the reference panel. Finally, there were 31 dogs from 14 breeds within our test samples with no previously assigned clade (**Supplemental Table S4**). Since most of these breeds are either European or have recent known European ancestry, they can likely be assigned to previously identified clades. Together, this data suggests the Plassais et al. (2019) dataset likely represents many of the same haplotypes found in the reference panel used for imputation, and that the test samples represent a mixture of shared and unique breeds, appropriate for determining the impact of imputation on low-pass sequencing.

**Table 1:** Clade representation of reference and test datasets

| Clades | Parker et al. (2017)<sup>1</sup> |  | Plassais et al. (2019)<sup>2</sup> |  | Test Samples |  |
| --- | --- | --- | --- | --- | --- | --- |
|  | Samples | Breeds | Samples | Breeds | Samples | Breeds |
| Alpine | 26 | 3 | 20 | 4 | 5 | 3 |
| American Terrier | 16 | 3 | 1 | 1 | 4 | 2 |
| American Toy | 20 | 2 | 5 | 2 | 0 | 0 |
| Asian Spitz | 83 | 9 | 25 | 9 | 1 | 1 |
| Asian Toy | 44 | 5 | 2 | 2 | 2 | 2 |
| Continental Herder | 44 | 5 | 25 | 4 | 2 | 2 |
| Drover | 34 | 4 | 15 | 4 | 0 | 0 |
| European Mastiff | 139 | 16 | 24 | 11 | 7 | 5 |
| Hungarian | 9 | 2 | 0 | 0 | 3 | 1 |
| Mediterranean | 98 | 14 | 12 | 6 | 7 | 3 |
| New World | 45 | 7 | 24 | 7 | 3 | 2 |
| Nordic Spitz | 64 | 5 | 13 | 10 | 7 | 4 |
| Pinscher | 12 | 2 | 0 | 0 | 1 | 1 |
| Pointer Setter | 88 | 12 | 18 | 10 | 5 | 3 |
| Poodle | 72 | 8 | 20 | 6 | 3 | 2 |
| Retriever | 66 | 7 | 48 | 7 | 3 | 2 |
| Scent Hound | 71 | 8 | 14 | 6 | 1 | 1 |
| Schnauzer | 20 | 2 | 5 | 2 | 0 | 0 |
| Small Spitz | 14 | 2 | 1 | 1 | 1 | 1 |
| Spaniel | 44 | 5 | 14 | 5 | 2 | 1 |
| Terrier | 140 | 18 | 100 | 15 | 7 | 6 |
| Toy Spitz | 32 | 4 | 2 | 2 | 2 | 2 |
| UK Rural | 145 | 16 | 42 | 12 | 0 | 0 |
| Unplaced*<sup>3</sup> | 20 | 2 | 7 | 4 | 0 | 0 |
| Unknown breed*<sup>4</sup> | 0 | - | 13 | - | 0 | - |
| Village Dogs*<sup>5</sup> | 0 | - | 69 | - | 0 | - |
| Mix Breed*<sup>6</sup> | 0 | - | 6 | - | 0 | - |
| Wild Canids*<sup>7</sup> | 9 | 2 | 29 | 2 | 0 | 0 |
| No Clade Info*<sup>8</sup> | 0 | 0 | 0 | 0 | 31 | 14 |

<sup>1</sup> Analysis that initially defined breed clade membership
<sup>2</sup> Dogs in the reference panel that are also included in Plassais et al. (2019)
<sup>3</sup> Dog breeds that formed their own branch in previous phylogenetic analysis and were not a member of a clade
<sup>4</sup> Dogs with no corresponding breed information
<sup>5</sup> Non-breed dogs sampled from 14 distinct geographic regions
<sup>6</sup> Dogs with mixed breed ancestry
<sup>7</sup> Group consists of grey wolves and golden jackals and breed label is used to differentiate these two different species
<sup>8</sup> Breeds not included in previous phylogenetic analyses and therefore not assigned clade membership
\* Groups of samples that either do not form a monophyletic clade or have not been included in previous phylogenetic analyses

### Downsampling and imputation

The 97 test samples were each downsampled to a coverage level of 1x and underwent imputation using loimpute as part of the Gencove, Inc. platform (Wasik et al. 2021). The average read coverage of WGS variants and imputed variants was 17.5x and 1.06x, respectively (**Supplemental Table S5**). A single sample, Pointer06, had a mean coverage of 1.67x, an outlier compared to next highest coverage dog which was 1.30x, suggesting Pointer06’s original coverage levels were incorrectly estimated. However, since variation in coverage level is a potential outcome of low-pass sequencing, Pointer06 was retained for further analyses. Imputation returned 53,649,170 million (M) variant sites, consisting of 35,875,925 SNV and 17,773,245 indel sites. Most sites were homozygous for the reference allele across all samples and were therefore removed from the analysis, leaving a total of 14,845,499 SNVs and 7,946,973 indels. Alternatively, genotype calling of high coverage WGS data for the test samples resulted in 18,476,517 SNVs and 12,831,692 indels.

To analyze the accuracy of imputation, we identified variants shared between the following groups: low-pass imputation, high coverage WGS, and the Plassais et al. (2019) dataset. The majority of SNVs, 13,943,807, were shared between all three variant groups, which represented 93.9% of all imputed SNVs. Similarly, the majority of indels, 5,333,851, were also shared between the three variant groups. However, shared indels represented only 67.1% of all imputed indels, a smaller fraction than observed with SNVs. Importantly, 99.6% of all imputed SNV sites and 88.6% of indel sites were present within the Plassais et al. (2019) dataset, indicating its utility as a stand in for the reference panel (**Fig 2A**).

**Fig 2:**
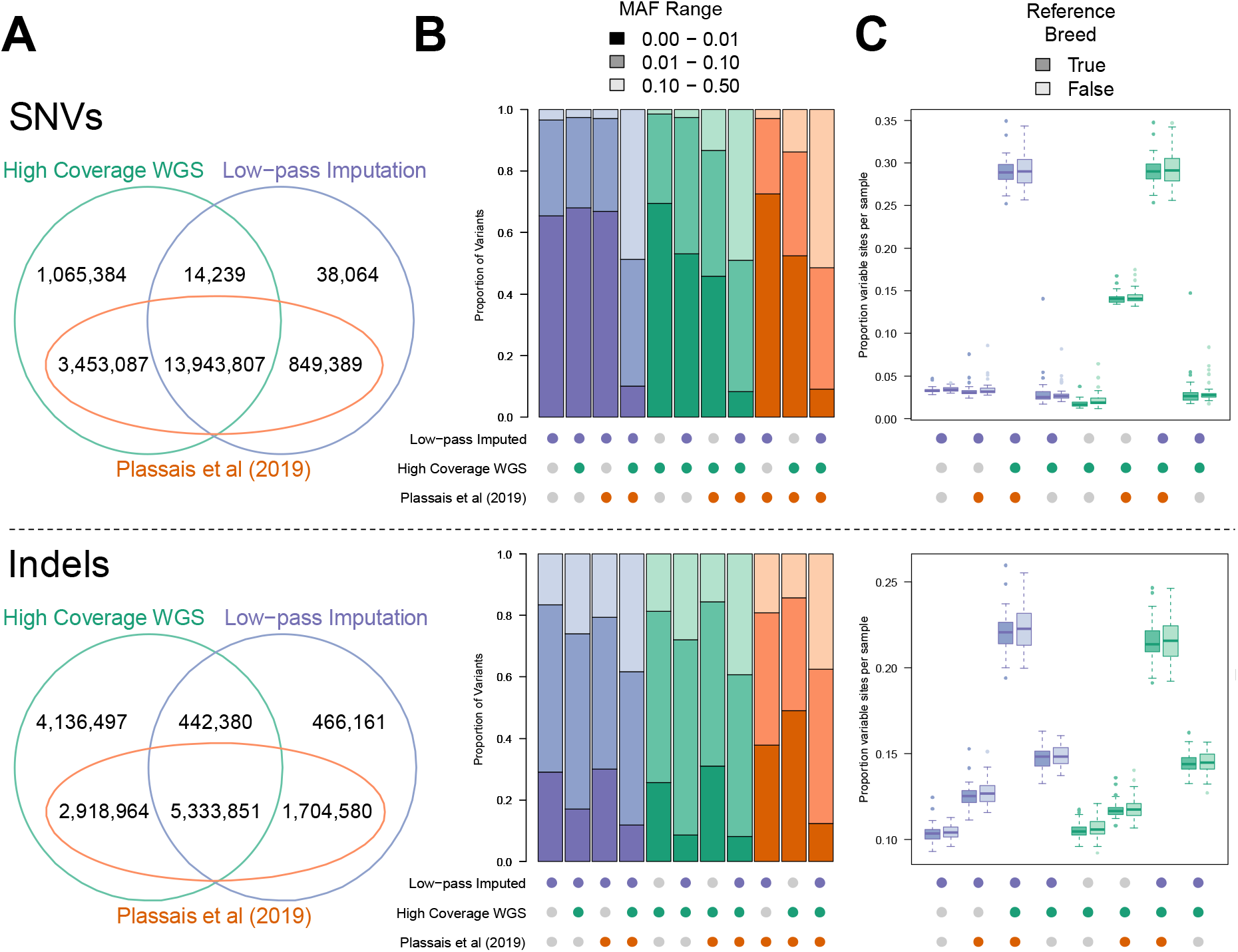
Genomic variant positions and their corresponding alleles are consistent across datasets. **A)** Venn diagrams for SNVs and indels showing variants unique and shared across datasets. Datasets include the high coverage WGS variant sites and low-pass imputed variant sites found across the 97 test samples and variants discovered in Plassais et al. (2019). Variants were identified as shared across datasets if the variant position, reference allele, and alternate allele were identical. **B)** MAF distribution of each variant group from A. Variant groups are indicated by colored circles beneath the bar chart. Groups contain variants which are the intersect between the colored circles and the set difference between the grey circles. The colors of each bar indicate the dataset used to calculate the MAF distribution and the shading level indicates the relevant MAF range. **C)** Sites per sample in each variant group, where variant groups are presented as in B. Sites per sample are measured as the proportion of total sites within the relevant variant group that contain a non-reference allele for a particular sample. Samples have also been divided into two groups based on whether the respective breed also belongs to the Plassais et al. (2019) dataset and is therefore likely used in the imputation reference panel.

Although there were high levels of agreement between the low-pass imputation and the high coverage WGS variant groups, many sites were specific to only one variant group. Since these sites were removed from our downstream analysis, we measured their MAF distributions to determine the extent of their potential impact on imputation accuracy measurements. Variant group specific sites usually had MAFs < 0.01, whereas variants shared across all groups usually had higher MAFs (**Fig 2B**). This indicated that high coverage WGS specific sites were likely individual specific sites, and absent from the reference panel used for imputation. Although a large number of these sites are also found in the Plassais et al. (2019) dataset, they may belong to breeds present in the test samples that were not present in the imputation reference panel. Low-pass imputation specific sites are due to imputation errors, as these sites are imputed as variable, but are actually genotyped as homozygous for the reference allele across all high coverage WGS samples. Since these variants tend to present as rare alleles (MAF < 0.01) in the imputed data, they are easily filtered out by MAF cutoffs typically used in association analyses and, thus, have only a minor impact on analysis.

To determine whether breed composition of the reference panel has an impact on variant imputation within out test samples, we counted the number of variant sites per individual for each combination of the variant groups and compared the results between reference panel breeds and non-reference panel breeds. Results showed that each carried a similar number of variant sites for both SNVs and indels, indicating that the breed composition of the current reference panel has little impact on variant imputation (**Fig 2C**). Ultimately, the total number of variant sites per samples varied according to MAF distributions, as shown in **Fig 2B**.

### Filtering strategies to optimize accuracy

Filtering strategies were optimized by analyzing the relationships between imputation accuracy, genotyping confidence as determined by max genotype probability (GP), and low confidence genotype thresholds (Methods) (**Supplemental Fig S1**). Receiver operator characteristic (ROC) curves were calculated for seven GP thresholds and eight low confidence genotype thresholds (**Supplemental Fig S2**). Each curve followed a similar trajectory. However, at a specific GP threshold, low confidence genotype thresholds lead to different outcomes for true positive rates (TPRs) and false positive rates (FPRs). For example, at GP > 0.7, a low confidence genotype threshold of two is required for a TPR > 0.8 and a FPR > 0.4 (**Fig 3A**). By comparison, to meet those same criteria at a threshold of GP > 0.9, a low confidence genotype threshold of four is required (**Fig 3B**). These results indicate that at higher GP thresholds, a greater level of robustness is achieved when selecting a low confidence genotype threshold, as small changes in threshold values do not lead to large changes in the number of variants removed.

**Fig 3:**
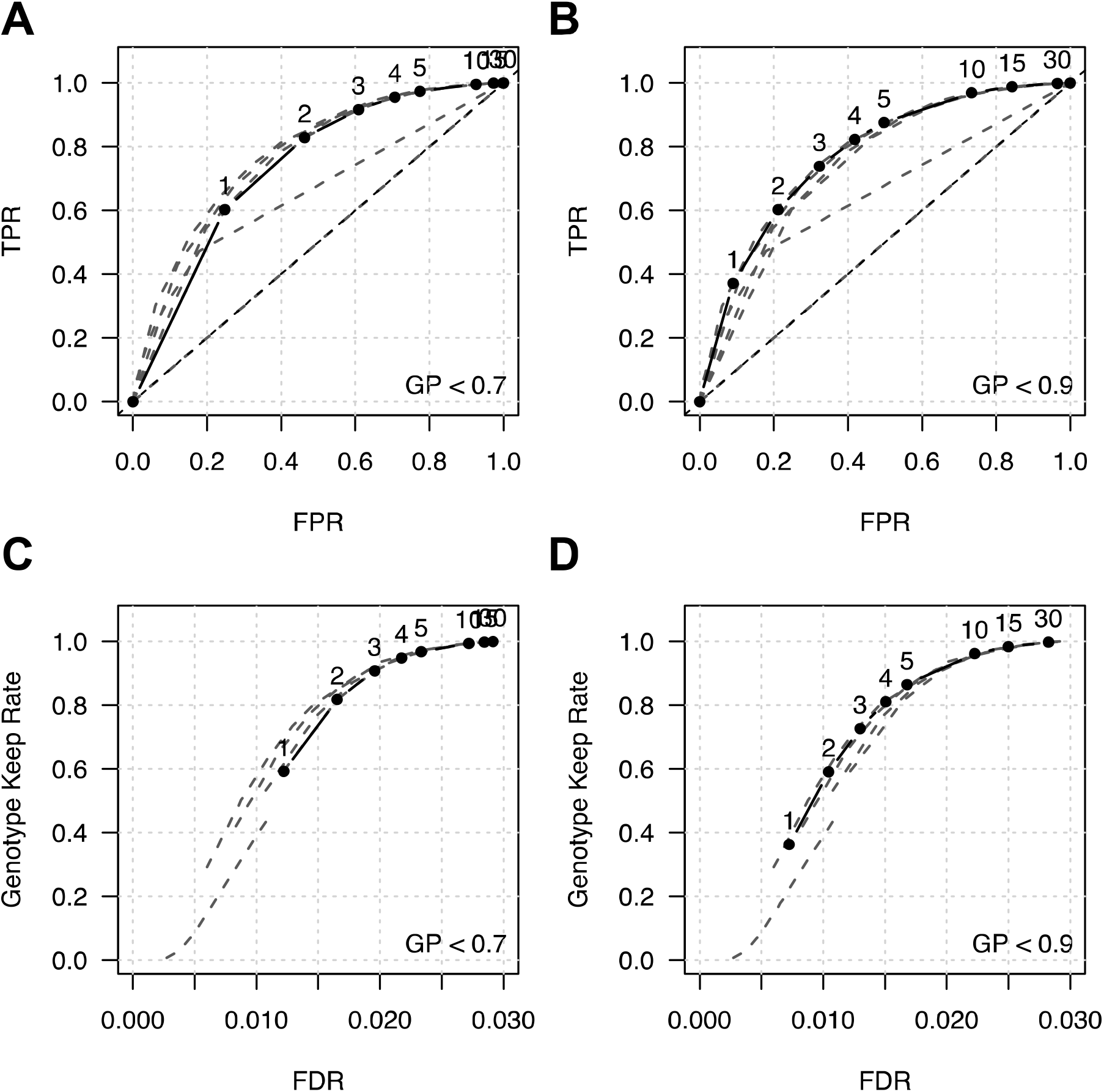
Performance of filtering strategies for reducing imputation errors. **A)** ROC curve, where genotypes with GP < 0.7 are identified as low confidence. Numbers above each point represent low confidence rate thresholds for removing sites. Sites with a total number of low confidence genotypes greater than or equal to the threshold are removed. Grey dashed lines represent ROC curves for other confidence threshold values. **B)** ROC curve for confidence threshold set at GP < 0.9. **C)** The proportion of variants remaining after filtering genotypes at GP < 0.7 and the corresponding FDR. As in C, the numbers above each point represent low confidence rate threshold values and grey dashed lines represent curves for other confidence thresholds. **D)** Proportion of variants remaining and their corresponding FDR after filtering at GP < 0.9.

False discovery rate (FDR) and keep rate were also analyzed for the thresholds mentioned above (Methods) (**Supplemental Fig S3**). This was done to assess how much data was lost by filtering and to assess the number of imputation errors that remain within the dataset after filtering. Similar to results shown in ROC curves, a higher GP threshold provided a higher level of robustness for selecting a low confidence genotype threshold. For example, at GP < 0.7, a low confidence genotype threshold of two leads to a keep rate > 0.8 and an approximate FDR of 0.015 (**Fig 3C**). To achieve a similar keep rate and FDR at GP < 0.9, a low confidence genotype threshold of four is required (**Fig 3D**). These results show that higher GP thresholds are more suitable for filtering sites based on the number of low confidence genotypes. This strategy allows for fine tuning of filtered results without a loss in accuracy. For the remainder of our analysis, we used a confidence filter of GP < 0.9 and a low confidence genotype threshold of four, which roughly corresponds to a genotyping error rate of 5%.

### Minor allele frequency impacts imputation accuracy

Imputation often performs poorly for rare alleles where statistical support is lacking. To determine the impact of low MAFs on low-pass imputation in dogs, we analyzed imputation accuracies using imputation quality score (IQS), a statistic that controls for allele frequencies by taking chance agreement into account (Lin et al. 2010). In addition, population MAF estimates were sourced from two different datasets, the imputed genotypes and the Plassais et al. (2019) dataset. These were chosen as each is available for similar low-pass imputation analyses.

Both sources of MAFs expressed similar IQSs, where the largest differences are due to whether imputed data was filtered. At MAFs > 0.1, the IQS of the unfiltered genotypes plateaued at approximately 0.91, whereas filtered genotypes plateaued at approximately 0.95 (**Fig 4A**). At MAFs < 0.1, the most accurate method for analyzing imputed genotypes is to use MAFs generated from imputed genotypes. At a MAF of 0.05, only the filtered genotypes that were partitioned according to imputed MAFs had an IQS > 0.9, providing the most accurate results for low MAF genotypes.

**Fig 4:**
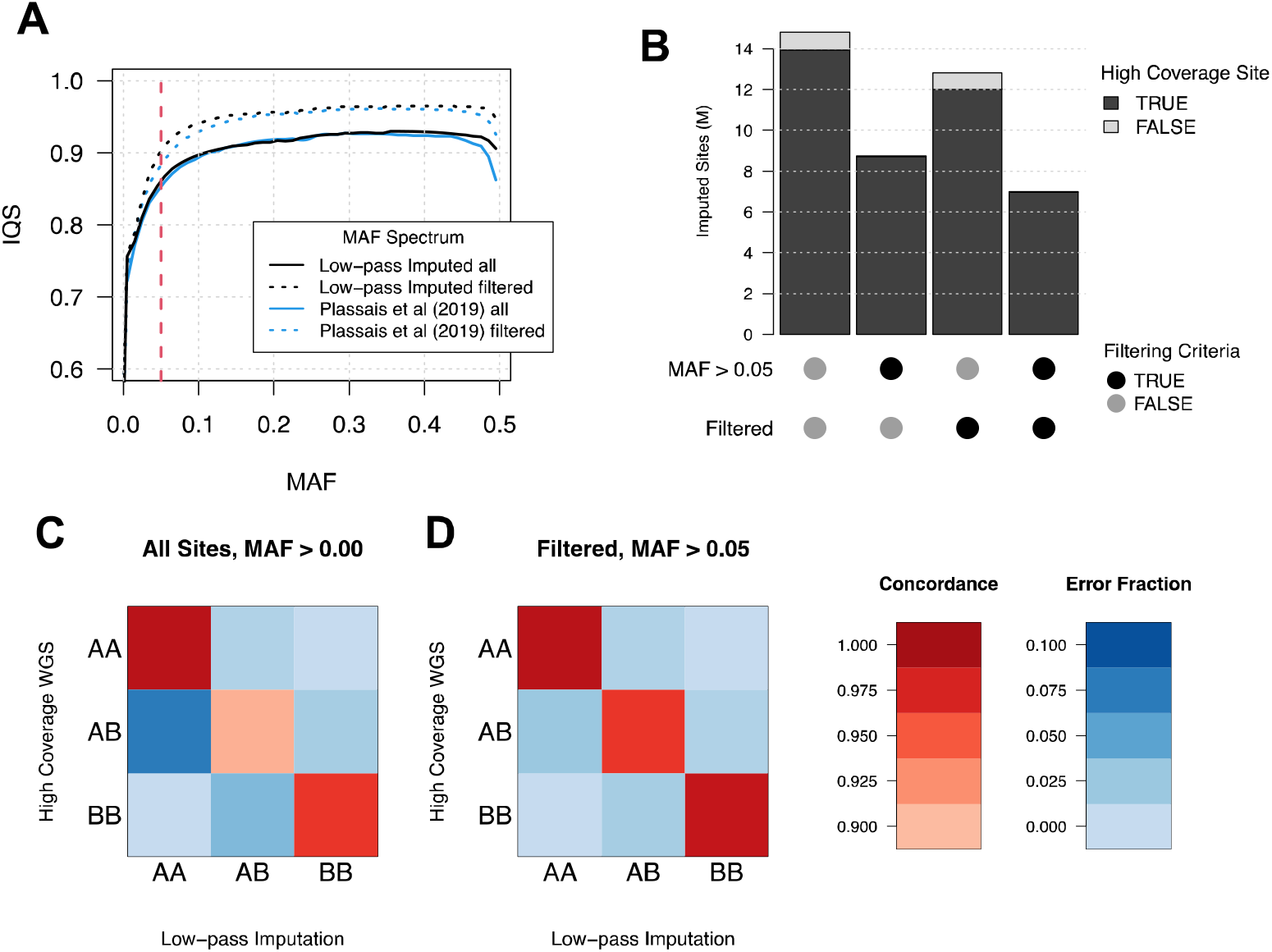
Imputation accuracy according to minor allele frequency and genotype. **A)** Imputation accuracy according to imputed and Plassais et al. (2019) MAFs for all sites and quality filtered sites. Imputation accuracy is measured as mean imputation quality score (IQS), an imputation accuracy statistic that accounts for the probability an allele is correctly imputed by chance. The red dotted line indicates a MAF of 0.05. **B)** The number of sites remaining after filtering for MAF > 0.05 and for low confidence genotypes < 5% as indicated by the “Filtered” label. Bar colors represent imputed sites that were either found or missing from the high coverage WGS dataset. **C)** Concordance and error rates for all genotypes, expressed as a fraction of the total number of high coverage WGS genotypes. **D)** Concordance and error rates for genotypes in sites with < 5% low confidence genotypes and MAFs > 0.05. Rates are expressed as a fraction of the number of high-coverage WGS genotypes that meet the corresponding filtering criteria.

Next, we analyzed the impact of MAF and filtering on imputation accuracy for different genotypes. Imputation accuracy was poorest for heterozygous genotypes, especially at low minor allele frequencies, indicating that heterozygous genotypes were least likely to be correctly imputed. Conversely, homozygous reference imputation was most accurate at lower MAFs, which is likely due to increased chance agreement between the high number of reference genotypes (**Supplemental Fig. S4**). In addition, non-reference concordance, mean *r*^2^, and IQSs were measured across chromosome 38 so that results can be compared to other analyses that use a different accuracy measurement (**Supplemental Fig. S5**). All three measures show higher levels of imputation accuracy at higher MAFs and a clear improvement for accuracy measurements for filtered genotypes. Together these results demonstrate that removing sites with a MAF < 5% retains a highly accurate set of genotypes, whereas filtering on GP values increases overall imputation accuracy. In addition, we show imputation errors are more likely for heterozygous sites, and MAF estimates derived from imputed genotypes are suitable for filtering by MAF.

An important consideration when filtering variants is the total number of sites remaining. Removing variants with a MAF < 0.05 and filtering out variants according to GP values results in a total of 7M remaining sites. Most sites are removed due to the MAF cutoff rather than GP filtering. In addition, almost all sites remaining after filtering correspond to a site found in the high coverage WGS dataset, highlighting the quality and accuracy of the final dataset (**Fig 4B**). An additional impact from filtering is the reduced variability between genotype specific imputation errors and concordance rates. For example, prior to filtering the overall concordance rate for high coverage WGS heterozygous genotypes with low-pass imputed genotypes was 0.908 with 0.072 of those imputed as homozygous reference, and 0.020 imputed as homozygous alternate (**Fig 4C**). Conversely, after filtering, 0.964 of these genotypes were imputed correctly, with approximately 0.023 genotypes imputed as homozygous reference and 0.013 genotypes as homozygous alternate (**Fig 4D**). Therefore, after filtering according to GPs and MAFs, overall concordance rates are increased relative to unfiltered genotypes and imputation errors are more evenly spread across the other genotypes.

To determine whether the breed composition of the refence panel used for imputation impacts imputation accuracy, we measured non-reference concordance according to breed and MAF. Overall, breeds whose members showed the highest levels of imputation accuracy prior to filtering, such as the Entlebucher sennenhund, Belgian malinois, Bernese mountain dog, Portuguese water dog, and Belgian tervuren, also had members within the imputation reference panel (**Fig 5A**) (**Supplemental Table S6**). Importantly, four of these five breeds were among the top 20 most highly represented breeds within the reference panel, with each containing at least nine members each within the reference panel (**Fig 1C**). Alternatively, breeds with the lowest levels of imputation accuracy usually did not contain members with the reference panel (**Fig 5A**). Moreover, breeds with low imputation accuracy that did have members in the reference panel, such as the Samoyed and Keeshond, had poorer representation than high imputation accuracy breeds, with both breeds containing only three members each within the reference panel (**Supplemental Table S2**). Furthermore, imputation accuracy rates of reference panel breed members were consistently higher than non-reference panel breed members across all MAF ranges (**Fig 5B**). These results indicate the importance of breed representation for improving imputation accuracy.

**Fig 5:**
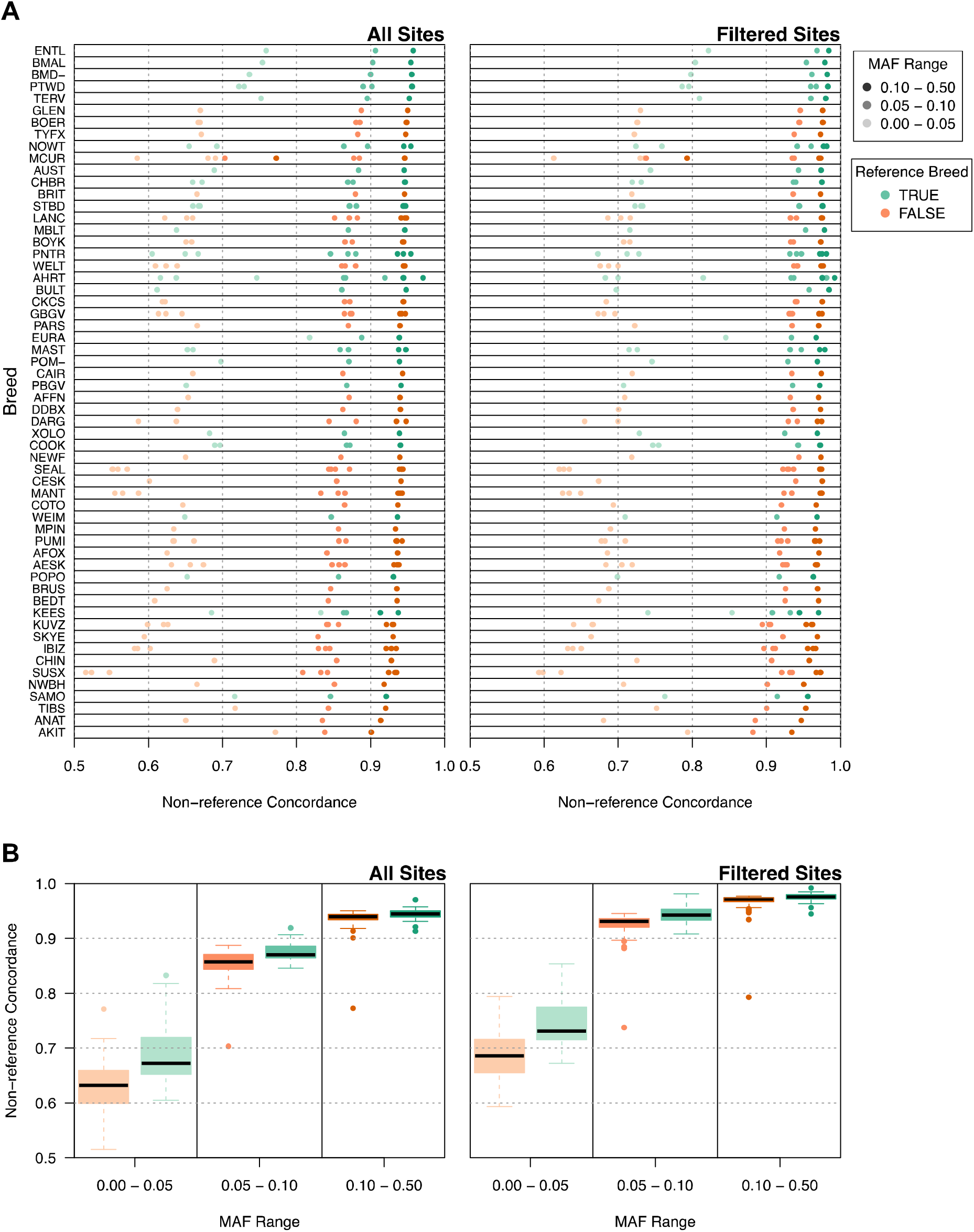
Imputation accuracy of dog breeds. **A)** Individual dog breed imputation accuracy. Dog breeds are displayed on the Y axis with imputation accuracy on the X axis as non-reference concordance. Accuracy rates are displayed for all sites (left) and sites that remain after quality filtering (right). The shading of each data point indicates imputation accuracy of SNVs within a specific MAF range. Green data points show breeds identified in the reference panel, while orange points show breeds not found in the reference panel. Breeds are ranked according to their median imputation accuracy for all sites. Imputation accuracies are displayed for each member of the breed. **B)** Imputation accuracy of reference and non-reference breeds according to MAF.

### Imputation errors reduce statistical power

After characterizing genotyping errors introduced by imputation, we simulated the impact of these errors on case-control association tests to determine best practices for study designs involving low-pass imputed genotypes. Given a specific genotype detected by high coverage WGS within a specific 0.01 population MAF interval, imputation errors were characterized as the probability of imputing either a homozygous reference, heterozygous, or homozygous alternate genotype. Overall, the imputation process led to decreased significance levels, suggesting that imputation errors may cause statistical significance to be lost for certain experimental configurations (**Fig 6A**). To investigate the loss of statistical significance in the context of study design we performed a power analysis with a focus on the number of samples required to reach sufficient power at 0.8 (**Fig 6B**). Specifically, we tested 21 case and control population MAF combinations over three different case-control ratios and used the MAF of the entire population to simulate imputation errors (**Fig 6C**). Case and control genotypes were based on Hardy-Weinberg equilibrium and were calculated using each population’s MAF (Methods). Results showed that experimental configurations for which the difference between case and control was smallest required the largest sample sizes. The required samples sizes were also higher for scenarios with non 1:1 case-control ratios, indicating that asymmetric population sizes increase the impact of imputation errors.

**Fig 6:**
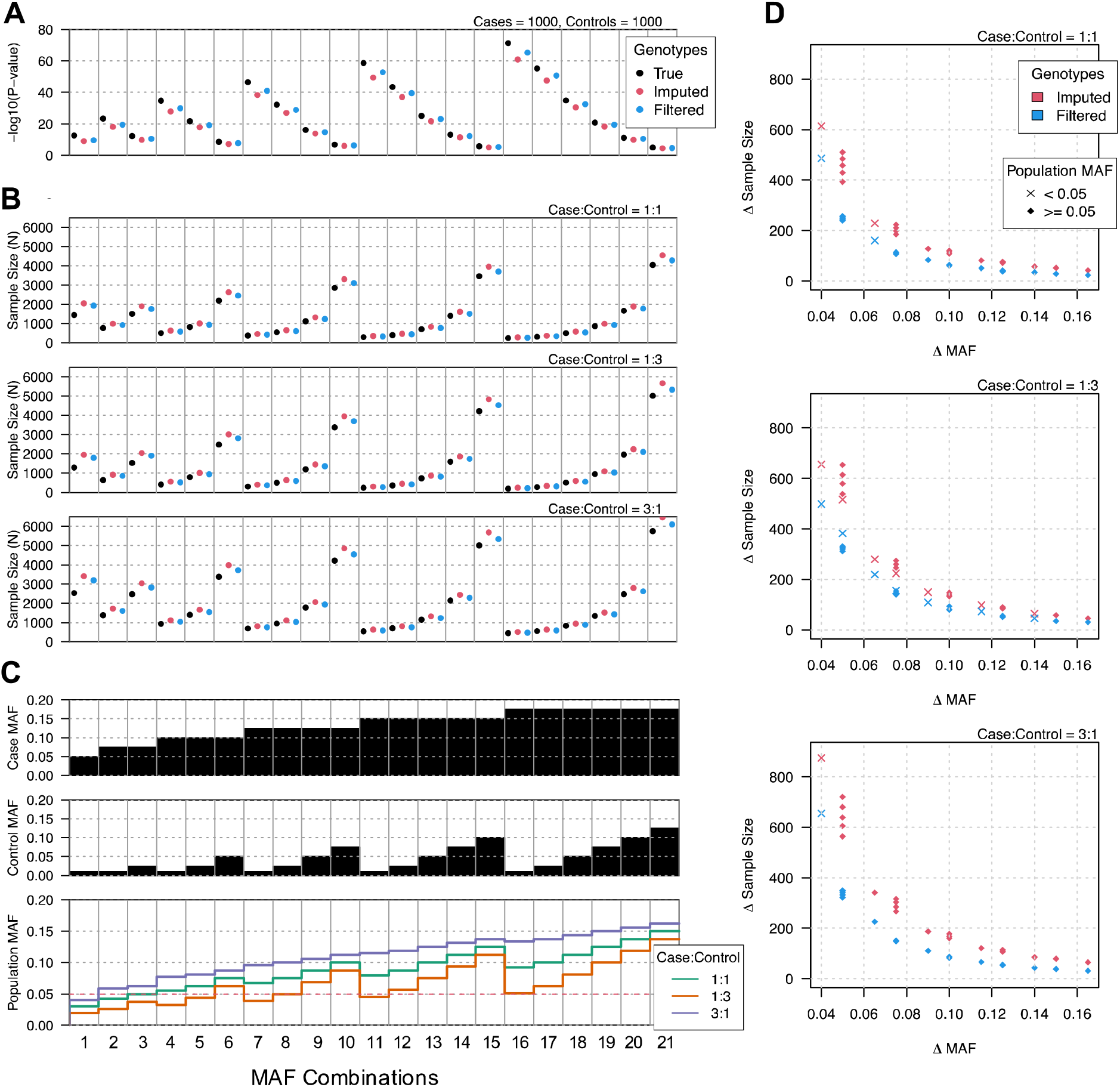
Impact of imputation errors on case-control GWAS. **A)** Significance of case-control GWAS at multiple MAFs. The true genotypes are represented by black circles, where the frequency of heterozygous and homozygous variants follow Hardy–Weinberg equilibrium. Red circles represent the outcomes of significance testing on imputed genotypes, while blue circles represent outcomes after filtering imputed genotypes. Note, decreases in significance were due to estimates of errors introduced during the process of imputation. Imputation errors were modeled according to the probability of a given genotype being imputed as any other genotype at any stated MAF. **B)** Power analysis of significance testing for case-control GWAS of true and imputed genotypes. Y axis shows required samples size to reach a statistical power of 0.80. Each individual plot shows different case-control ratios. Power was calculated for a 2×2 chi-square test for significance level 5×10^−8^, where effect size was calculated as Cohen’s w. **C)** Case and control MAFs used for each significance test analysis and the combined population allele frequency for each case and control configuration. **D)** Additional samples required to reach sufficient power for imputed genotypes. Delta sample size is the difference between required sample sizes for true genotypes and imputed or quality filtered imputed genotypes. Delta MAF is the difference in MAFs between cases and controls. Delta MAF is proportional to effect size.

Importantly, the increased imputation accuracy afforded from quality filtered data translated to reduced sample size requirements for achieving sufficient power (**Fig 6B**). Also, the difference in MAFs between cases and controls was strongly predictive. MAF differences of 0.05 required > 500 additional samples for unfiltered imputed genotypes to achieve the same power as the true genotypes, while MAF differences of 0.1 required approximately 100 additional samples (**Fig 6C**). Similar to the higher number of total samples required for asymmetric populations, sample requirements for imputed data were also increased. Finally, the additional samples required for quality filtered imputed data is slightly greater than 50% of the additional samples required for unfiltered data (**Fig 6C**). Together, these results indicate the importance of considering imputation error in the role of study design. Importantly, the impact from imputation errors is inversely proportional to the effect size of the association.

## Discussion

Genotype imputation is a valuable tool for filling in missing genotypes and improving power to detect genome wide associations. It also provides an opportunity to combine samples genotyped on different platforms into a single analysis (Marchini et al. 2007; Uh et al. 2012; Hayward et al. 2019). Ultimately, this increases the amount of available data by allowing datasets to be reused in larger studies (Ho and Lange 2010; Zhuang et al. 2010). However, imputation techniques have gone a step further and now facilitate genotyping of poor quality or low-pass sequencing data, which often lack sufficient coverage for genotyping software to assign confident calls. Imputation reference panels and algorithms provide the additional statistical support required to assign genotypes to individual samples (Rubinacci et al. 2021; Wasik et al. 2021). However, since the samples used to impute genotypes are different from those that undergo low-pass WGS, imputation has the potential to introduce genotyping errors and biases. Therefore, we investigated the impact of imputation on genotyping in dogs, an important genetic system whose unique population structure, defined by strict breeding programs over hundreds of years (Ostrander et al. 2017), may uniquely influence imputation accuracy. To characterize imputation errors and provide best practices for analyzing low-pass imputation data in dogs, we tested imputation for 97 high converge WGS dog samples with approximately 1x coverage per sample. Imputation errors were detected by comparing the imputed genotypes to genotypes determined using high coverage WGS data. We analyzed these errors in the context of genotype quality filtering, imputation error biases, the role of MAFs, and their impact on case-control analyses.

Our analysis of case-control association tests of imputed genotypes provides the necessary information to outline best practice guidelines for working with low-pass imputed genotypes. We show herein that the most important factor to consider is the expected allele frequencies in both cases and controls for any potentially associated markers. This is important, as the most common error observed was associated with imputing a heterozygous genotype as homozygous reference, leading to an overall reduction in observed effect size and therefore requiring more samples to reach sufficient power. Importantly, these reduced associations were most pronounced when the allele frequency difference between cases and controls was small (≤ 0.05). Therefore, best practices in study design for low-pass imputation are to investigate genetic associations with medium to large effect sizes. Alternatively, if effect sizes are likely small, investigators need to consider increased sample sizes, balance between case and control populations, and the role of quality filtering for improving overall accuracy.

Quality filtering imputed genotypes is achieved by removing sites where ≥ 5% of genotypes have maximum genotype probabilities (GP) < 0.9. This threshold was chosen as it is robust to small shifts in genotype failure rate, which causes only a small change in the total number of sites removed. The genotype failure rate threshold of 5% is well-calibrated to optimally reduce imputation errors at a GP threshold of 0.9. We demonstrate that quality filtering is able to improve IQSs by a value of ~0.04 and reduce required increases in sample sizes for sufficient power by as much as 50%. Essentially, these improvements are achieved by removing ~ 20% of sites that together share a disproportionate number of the imputation errors. In addition, removing sites with MAFs < 0.05 removes those with comparatively lower imputation accuracies and sites that aren’t found in high coverage WGS datasets.

After both quality and MAF filtering ~ 7M SNVs remain, with SNVs found approximately every 360 bp. Despite removing the majority of SNV markers through filtering sufficient numbers of SNVs remain for association analyses. Typically, association studies within breeds demarcate regions on the order of one Mb (Sutter et al. 2004; Lindblad-Toh et al. 2005; Vaysse et al. 2011; Karlsson et al. 2013), whereas across breeds, the scale of LD is approximately 10 – 100 kb (Karlsson et al. 2007; Parker et al. 2017). In addition, currently available canine DNA genotyping arrays contain just over 710,000 markers (Axiom^™^ Canine HD Array), one tenth the total number of markers available from low-pass imputation after quality and MAF filtering. Therefore, the benefit of increased genotyping accuracy using filtering likely exceeds the cost incurred from reduced marker density.

An additional consideration in performing GWASs using imputed data with small effect sizes is the required increase of both the case and control populations to reach sufficient power. For filtered genotypes with population MAF differences of 5%, approximately 250 additional samples with equal proportions of cases and controls are required to reach sufficient power. Importantly, as the MAF difference between cases and controls decreases, the required increase to sample sizes appears to grow exponentially. This is perhaps linked to increased rates of imputation errors at MAFs < 0.05.

Finally, for best practices study design, balance between case and control populations should also be addressed. The impact on power from unbalanced case and control populations is most prominent at low MAF differences between cases and controls. When the ratio between cases and controls favors either population, there is an overall loss of power compared to when the ratio is even. For example, at a case-control MAF difference of 5%, where the ratio of cases to controls is either 1:3 or 3:1, > 300 additional samples are required to reach sufficient power, whereas if the ratio is 1:1, only 250 additional samples would be required. This is because a lower number of total associated alleles, as opposed to proportion of alleles, within either cases or controls, increases the likelihood that the accumulation of imputation errors can cause statistical significance to be lost.

There are two strategies for developing reference panels, the first is to use a population with closely matched ancestry to that of the group under study, and the second is to use as many samples as possible. While not evaluated in the context of low-pass sequencing imputation, analysis of DNA arrays shows that reference panels matched to the population of interest outperform diverse reference panels of similar sizes (Mitt et al. 2017; Zhou et al. 2017; Bai et al. 2019; Yoo et al. 2019). This would suggest that larger refence panels are preferable as long as they contain sufficient representation of the study population. However, the addition of diverse samples to a reference panel can decrease imputation accuracy at low MAFs, where the magnitude of this effect varies according to the population being studied (Bai et al. 2019). These observed effects were for initial population-matched reference panels of ~100 samples, with additional diverse samples increasing the reference panels to over 860 samples (Bai et al. 2019). Other analyses compared references panels of > 1500 samples to the Haplotype Reference Consortium (HRC) reference panel (http://www.haplotype-reference-consortium.org/) (McCarthy et al. 2016; Mitt et al. 2017; Zhou et al. 2017; Yoo et al. 2019), which consists of 32,611 samples, indicating the potential for increased resolution in human studies compared to canine studies, which used a panel of just 676 samples from 91 breeds (Piras et al. 2020). Altogether, at MAFs > 0.05, human imputation studies conducted using DNA array genotypes show non-reference concordance rates > 97.5% and mean *r*^2^ values > 0.95 (Mitt et al. 2017; Zhou et al. 2017; Yoo et al. 2019). By comparison, prior to filtering non-reference concordance rates for low-pass sequence imputation in dogs were at ~ 95% and had mean *r*^2^ values between 0.90 and 0.94 (**Fig 4C**), highlighting the potential gains that can be achieved from improved canine reference panels.

An important initiative that may help address short comings in available high coverage WGS samples in dogs is the Dog 10K project which aims to achieve 10,000 modest-high coverage dog genomes representing an array of canine genetic diversity (Ostrander et al. 2019). Similar to the current dog reference panel, the initial phase of the Dog 10K project prioritizes collecting samples from as many modern breeds as possible. Alternatively, human data suggests that for imputation purposes it is better to use samples from the population of interest. Whether this same strategy is preferable in dogs is unknown.

Many mapping studies in dogs focus on traits that segregate across breeds, as breeds sharing recent common ancestry likely share the same genetic underpinnings for any given trait (Parker et al. 2017). While multiple breeds are often included within a single analysis, creating many breed-specific reference panels is not feasible. As large haplotype blocks are shared between many breeds and clades, perhaps a large reference panel representing a greater number of breeds could provide even higher levels of imputation accuracy than a breed-specific reference panel. This idea is supported by an array-based imputation analyses that tested the imputation accuracy for a group of poodles with three different refence panel configurations. Results showed a composite panel of poodles and non-poodles outperformed the poodle only and the non-poodle reference panels (Friedenberg and Meurs 2016). A key finding from our analysis, was that SNV discovery in individual dogs was similar between reference panel and non-reference panel breeds (**Fig 2C**), and while for imputation accuracy rates reference panel breeds scored highest, several non-reference panel breeds outperformed the majority of reference panel breeds (**Fig 5A**). This was likely due to the fact that many of breeds from the test samples belonged to clades represented in the reference panel. However, since the original VCF for the reference panel was unavailable, breeds were identified from matching IDs across databases and breed membership for 122 dogs could not be determined. Also, many of the samples in the reference panel were either village dogs or other canid species. In our test samples, of the 31 dogs from 14 breeds not previously associated with a clade (**Supplemental table S3**), five were terrier breeds that likely belong to the primary terrier clade and two were spaniel breeds that can be assigned to the spaniel clade. Since breeds of European origin are heavily represented in our refence panel, and most breeds with no clade assignment are of European ancestry, it is likely that many of the non-clade assigned test sample haplotypes are at least partially represented in the reference panel. Supporting this idea of partial haplotype representation providing accurate imputation in dogs is the observation of high levels of imputation accuracy shared between mixed and pure breeds (Hayward et al. 2019). As more high coverage WGS samples become available through initiatives such as Dog 10K, optimal reference panel designs can be constructed.

One additional improvement in imputation accuracy may derive from the choice of imputation algorithm. Currently, available tools for imputation of low-pass WGS data include STITCH (Davies et al. 2016), Beagle (Browning and Browning 2016), GeneImp (Spiliopoulou et al. 2017), GLIMPSE (Rubinacci et al. 2021), and loimpute (Wasik et al. 2021). Our analyses used loimpute, as it had already been implemented with a dog reference panel and used by the canine genomics community (Piras et al. 2020). However, if low-pass sequencing imputation approaches in dogs are going to improve, other algorithms need to be appropriately assessed with accuracy across a range of MAFs. Currently, human studies demonstrate that GLIMPSE outperforms the other algorithms in terms of both accuracy and required computations resources (Rubinacci et al. 2021). The largest differences were observed for variants with MAFs < 1%. For common alleles, imputation accuracy was similarly high between almost all algorithms.

Our work provides the first in-depth analysis of low-pass WGS and imputation in canine genomics and can act as a road map for analysis in other non-human species. By comparing genotypes imputed from downsampled reads to a high coverage truth set, we have been able to rigorously investigate the nature of imputation errors and their biases. We were able to optimize filtering strategies to improve accuracy rates and also demonstrate the impact imputation errors have on case-control GWAS. Our results inform a series of best practices guidelines and demonstrated the utility of this quickly evolving resource for future analyses. Altogether, widespread adoption of low-pass sequencing and imputation within the canine genomics field, together with investment in developing improved reference panels, will lead to more high-powered analyses and successful discovery of genotype-phenotype associations.

## Methods

### Sample selection

Samples were selected on the basis of whether they belonged to known breeds, were absent from the reference panel, and had mean sequence coverage levels > 15x. Non-publicly available samples were not used in the reference panel and therefore provide an accurate test of imputation performance. Gencove Inc. provided a list of sample IDs, most of which were matched to known samples within the Plassais et al. (2019) dataset, which was used to represent variant population frequencies. Breed names were based on annotated records and clade membership was based on previously published results (Parker et al. 2017). Breeds with no recorded clade membership were assigned to a clade based on their phenotype, historical information or phylogenetic clustering in Plassais et al. (2019).

### Variant calling and imputation

Sample reads were mapped to CanFam3.1 using BWA-mem (Li 2013). Variant calling was performed using GATK4 best practices (McKenna et al. 2010). Base quality score recalibration and duplicate marking was applied to each sample (DePristo et al. 2011; Van der Auwera et al. 2013), and haplotypecaller was used for variant discovery (Poplin et al. 2017). Average coverage was estimated using Samtools depth tool (Li et al. 2009). To simulate low-pass sequencing, BAM files were downsampled to approximately 1x coverage using the DownsampleSam tool from GATK4. To obtain the correct coverage level the parameter “-p” was set as the sample’s mean coverage divided by one. Downsampled BAMs were converted to fastq files using samtools “fastq” function and were uploaded to Gencove, Inc. using the Gencove command line interface (CLI). Imputation was performed using loimpute as part of Gencove’s imputation pipeline with the “Dog low-pass v2.0” configuration (Piras et al. 2020; Wasik et al. 2021). Imputed genotypes were received from Gencove as a VCF for each individual. Individual VCFs were split according to chromosome. Each sample’s genotypes and genotyping statistics were merged to create a single dataset for each chromosome that contained all individuals. This task was performed using the program extract_genotype_wg.R which was written in R and used the vcfR package (Team 2013; Knaus and Grunwald 2017).

### Assessing imputation accuracy

Imputation accuracy was assessed by comparing imputed genotypes to high coverage WGS genotypes. This was made possible by identifying sites shared across both datasets. Sites were considered shared if position, reference allele, and alternate allele were identical. Importantly, all multiallelic sites in all VCFs were split into biallelic states using the bcftools “norm” function with the “-m –” parameter to ensure all potential allelic combinations were matched (Li 2011). Sites where all samples were homozygous for the reference allele were removed from the analysis. A genotype for a given individual at a particular site was considered concordant if the imputed genotype was identical to the genotype determined using high coverage WGS. Imputation errors were those that were not identical between the high coverage WGS dataset and imputed dataset for a given individual at a given site. The total number of concordant genotypes per sample was calculated using the program “venn_filter_wg.R”. Once filtering thresholds were determined (below), imputation accuracies were determined according to MAF intervals of 0.01 using the program “gt_by_af.R”.

### Determining filtering thresholds

The imputation process provides genotype probabilities as a measure of confidence regarding whether a call is homozygous reference, heterozygous, or homozygous alternate. The max genotype probability (GP) is the level of confidence for the imputed genotype. We therefore investigated the relationship between GP and the proportion of concordant genotypes between our imputed and high coverage WGS datasets in order to determine optimal strategies for filtering imputed variants. Low confidence or failed genotypes were identified according to their GP values and variant sites were filtered out if the number of low confidence genotypes was above a given threshold. (**Supplemental Fig. S1A**). Filtering strategies were evaluated according to the number of remaining genotypes after filtering and the proportion of these genotypes that were concordant with the truth set. These values were further investigated in terms of the true positive rate (TPR), false positive rate (FPR), false discovery rate (FDR), and the number of genotypes kept after filtering (**Supplemental Fig. S1B**).

### Simulation of imputation errors

Imputation errors were simulated using probabilities derived from the observed fraction of any given genotype that was incorrectly imputed. Error probabilities were also grouped according to population MAFs. For example, for sites with MAFs between 0.03 and 0.04, 5% of heterozygous sites may be imputed as homozygous reference, whereas for sites with MAFs between 0.10 and 0.11, 3% of heterozygous sites may be imputed as homozygous reference. This strategy is used to capture the MAF impact on imputation accuracy and error rates. The imputed genotypes, *I*, were stored as a 1 x 3 matrix, consisting of the imputed genotype counts for homozygous reference, heterozygous, and homozygous alternate genotypes. The values for *I* were calculated as *I* = *GP*, where *G* is the counts for the true starting genotypes stored in a 1 x 3 matrix and *P* is a 3 x 3 matrix of the probabilities a true genotype is imputed as any other genotype. In *P*, rows represent the true starting genotype and columns represent the imputed genotypes. Importantly, the values used for *P* depend on the population MAF and whether the genotypes were filtered for quality. A total of 100 *P* matrices were defined, with one matrix for each 0.01 MAF interval between 0 and 0.5 for both quality filtered and unfiltered imputation error rates.

### Association analyses and statistical power calculations

Case-control association analyses were performed as a chi square test on a 2 × 2 contingency table, measuring the association between cases and the presence of the minor allele. Association tests were carried out on the true genotypes, imputed genotypes, and quality filtered imputed genotypes. Power was calculated using the “pwr” package in R (Champely 2020), which uses Cohen’s *w* to calculate effect size. Probabilities for the null hypothesis were calculated as if the minor allele was evenly distributed across both cases and controls. Significance levels were set at 5×10^−8^ and power was calculated across a variety of case and control MAFs for all population sizes between 100 and 7000. Required sample sizes for sufficiently powered analyses were identified as the lowest sample size that could achieve a power level of 0.8 or greater for a particular case-control analysis.

### Software and data analysis

All original software used here can be obtained from the following URL: https://github.com/NHGRI-dog-genome

## Data Access

Test samples used in this analysis are deposited under the BioProject accession PRJNA648123. Individual accessions for each sample are recorded in Supplemental Table S1.

## Acknowledgements

All authors acknowledge Gencove, Inc. for their assistance and guidance throughout the project. R.M.B., A.C.H., D.T.W., and E.A.O. were supported by the Intramural Program of the National Human Genome Research Institute at the National Institutes of Health. Support was also provided by the National Key R&D Program of China (2019YFA0707101), Key Research Program of Frontier Sciences of the CAS (ZDBS-LY-SM011), and Innovative Research Team (in Science and Technology) of Yunnan Province (202005AE160012). G.D.W. is supported by the Youth Innovation Promotion Association of CAS. We especially thank dog owners who have provided samples from their dogs for these and other studies.

## Supplemental Tables

Supplemental Table S1: Sample-specific metadata.

Supplemental Table S2: Breed composition of datasets.

Supplemental Table S3: Breed clade membership.

Supplemental Table S4: Breeds not included in previous phylogenetic analyses and have no clade membership.

Supplemental Table S5: Mean chromosome 38 coverage levels at variant sites.

Supplemental Table S6: Imputation accuracy rates for individual samples measured as non-reference concordance.

