## Supplemental Fig for "Best practices for analyzing imputed genotypes from low-pass sequencing in dogs"

Supplemental Figures

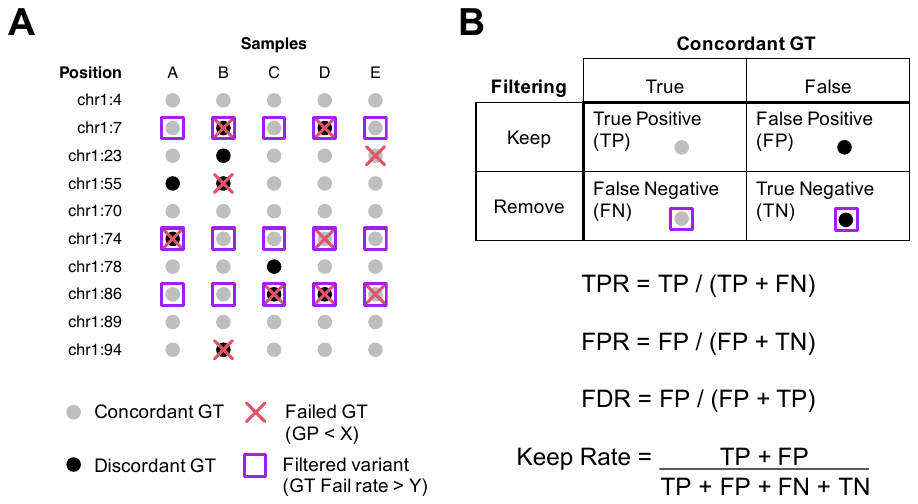

**Supplemental Figure S1: Filtering strategies for reducing imputation errors.** **A)** Schematic of imputed genotypes. Genotypes are represented as filled in circles, where black circles indicate discordant genotypes and grey circles indicate concordant genotypes. In this example the genotypes themselves, such as heterozygous and homozygous, are hidden as they are not relevant. Generally, genotype concordance between actual and imputed data remains unknown and other alternative metrics are used to filter out sites that likely contain an abundance of imputation errors. Here, max genotyping probability (GP) is used to assess genotyping confidence. GP below a certain threshold, X, identifies low confidence genotypes, which are marked with a red cross. Genomic positions that contain greater than a certain number of low confidence genotypes are filtered out as their low confidence genotyping rate is above the threshold Y. Here, sites with a low confidence genotyping rate > 20%, or 1 out of 5 samples are marked with purple squares. Ideally, sites removed by filtering are enriched for discordant genotypes. **B)** The statistics used to assess and compare filtering strategies. These include, true positive rare (TPR), false positive rate (FPR), false discovery rate (FDR), and keep rate, which is measured as the proportion of genotypes remaining after filtering.

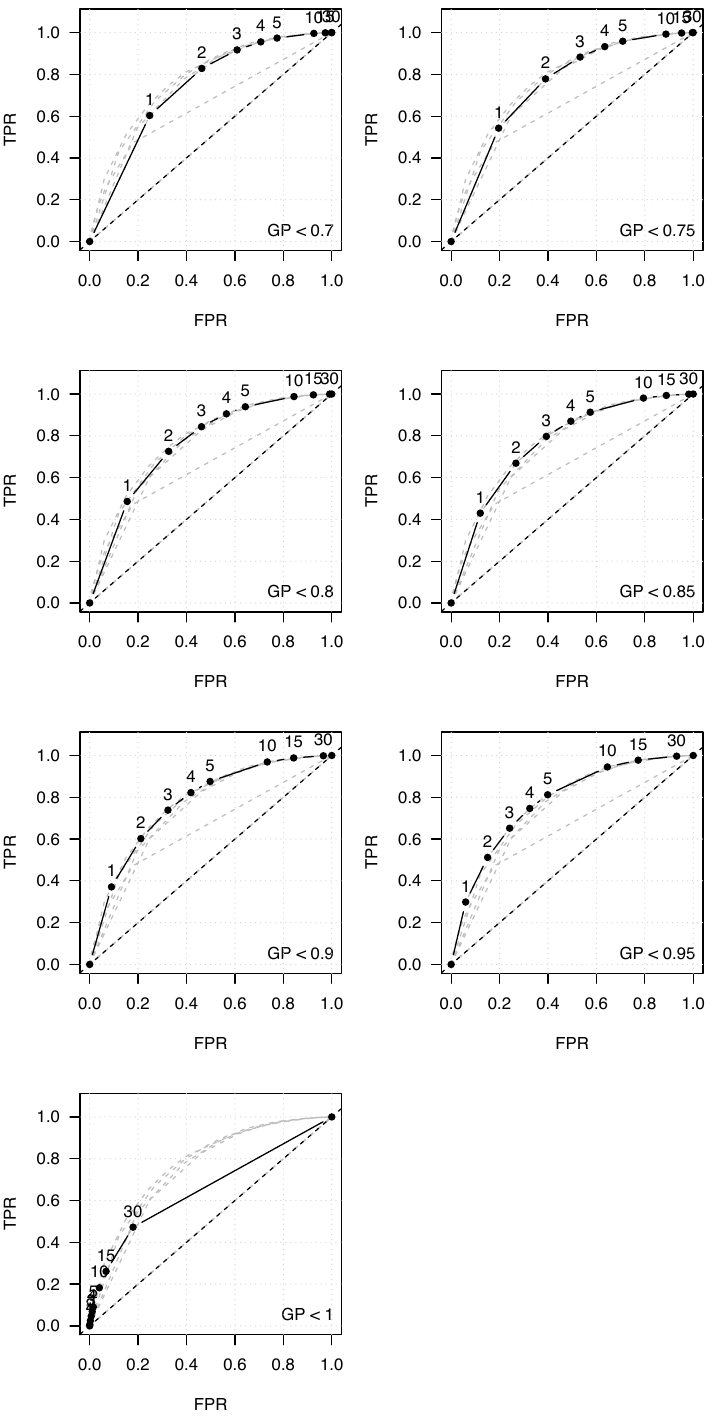
**Supplemental Figure S2: Receiver operator characteristic (ROC) curves at different GP thresholds.** Genotypes below GP thresholds shown in the lower right corner of each plot are designated as low confidence. Numbers above each point represent low confidence rate thresholds for removing sites. Sites with a total number of low confidence genotypes greater than or equal to the threshold are filtered out. Grey dashed lines represent ROC curves for other GP confidence threshold values.

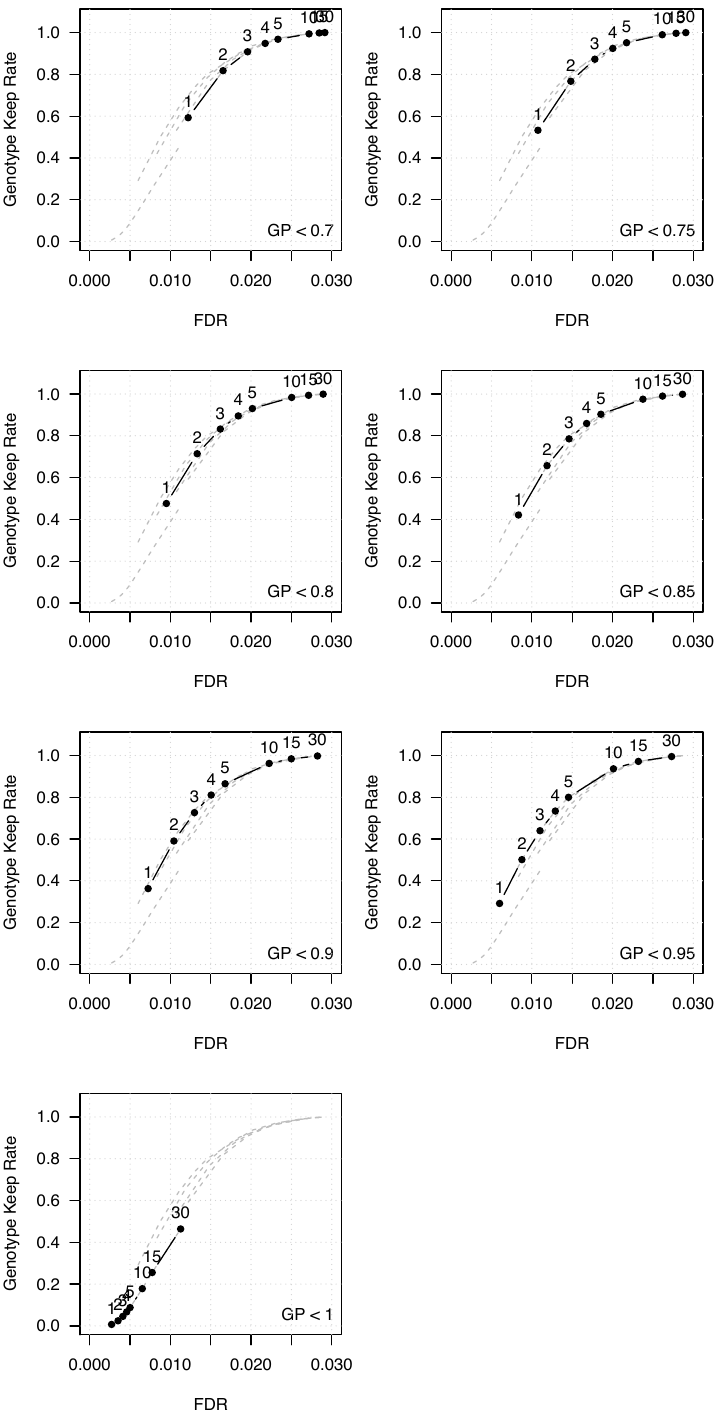
**Supplemental Figure S3: The proportion of variants remaining after filtering at different GP thresholds and the corresponding FDR.** Again, as in Supplemental Figure S2, genotypes below GP thresholds shown in the lower right corner of each plot are designated as low confidence. The numbers above each point represent low confidence rate threshold values and grey dashed lines represent curves for other GP confidence thresholds.

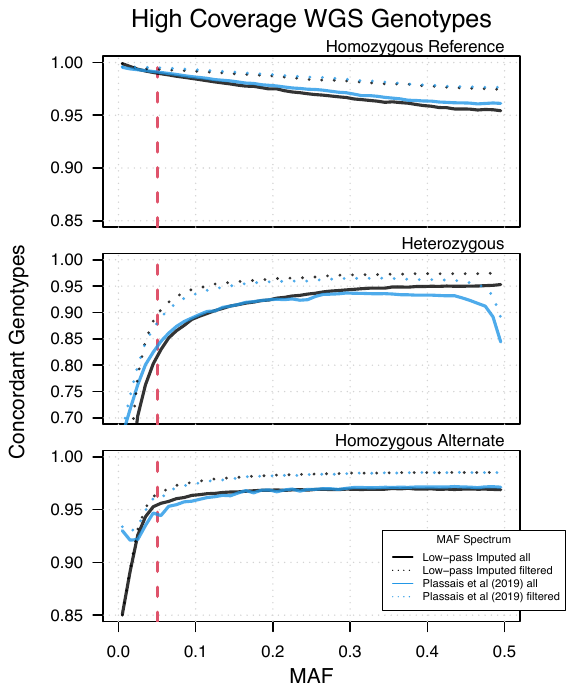

**Supplemental Figure S4: Imputation accuracy for each individual allele according to various MAF spectra.** Imputation accuracy is expressed as the proportion of high coverage WGS genotypes concordant with low-pass imputed genotypes.

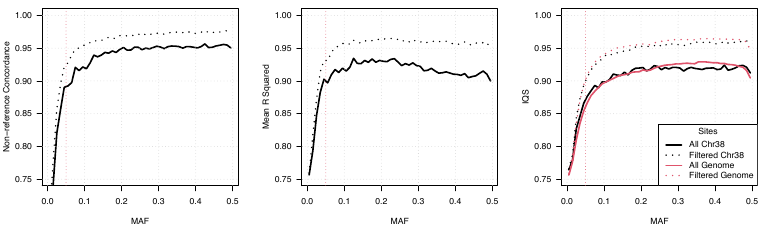

**Supplemental Figure S5: Comparison of various imputation accuracy measurements across chromosome 38.** Accuracy measurements including non-reference concordance, mean *r*^2^, and IQS. Whole genome measurements are included for IQS to show chromosome 38 is representative of the whole genome.
